# Opposing pupil responses to offered and anticipated reward values

**DOI:** 10.1101/298844

**Authors:** Tyler Cash-Padgett, Habiba Azab, Seng Bum Michael Yoo, Benjamin Y. Hayden

**Author notes:** Corresponding author: Tyler Cash-Padgett, Department of Neuroscience, University of Minnesota, Minneapolis, MN 55455.

## Abstract

Previous studies have shown that the pupils dilate more in anticipation of larger rewards. This finding raises the possibility of a more general association between reward amount and pupil size. We tested this idea by characterizing macaque pupil responses to offered rewards during evaluation and comparison in a binary choice task. To control attention, we made use of a design in which offers occurred in sequence. By looking at pupil responses after choice but before reward, we confirmed the previously observed positive association between pupil size and anticipated reward values. Surprisingly, however, we find that pupil size is negatively correlated with the value of offered gambles before choice, during both evaluation and comparison stages of the task. These results demonstrate a functional distinction between offered and anticipated rewards, and present evidence against a narrow version of the *simulation hypothesis*, the idea that we represent offers by reactivating states associated with anticipating them. They also suggest that pupil size is correlated with relative, not absolute, values of offers, suggestive of an accept-reject model of comparison.

## INTRODUCTION

The pupils systematically dilate and constrict in response to ongoing changes in mental state. Pupil diameter therefore provides a window into many important mental functions, ranging from attention (Hoeks and Levelt, 1993; van den Brink et al., 2016) and working memory (Kahneman and Beatty, 1966) to mental effort (Just et al., 2003; Varazzani et al., 2015) and surprise (Lavín et al., 2013; Preuschoff et al., 2011). Researchers have even used pupil size to gain insight into the mechanisms of subjective time perception (Suzuki et al., 2016), rate of learning (Nassar et al., 2012), and multi-sensory integration (Rigato et al., 2016), as well as decision-making (de Gee et al., 2014; Einhauser et al., 2010; Einhauser et al., 2008).

Previous research supports the idea that there is a positive relationship between reward magnitude and pupil size. Specifically, pupil size increases in anticipation of rewards and increases more in anticipation of larger primary rewards (Rudebeck et al., 2014). The positive relationship between pupil size and anticipated rewards is also observed in anticipation of conditioned reinforcers (Rudebeck et al., 2014; Varazzani et al., 2015). These results suggest that there may be a positive relationship between pupil size and reward amount that is observed for types of rewards other than anticipated ones.

We are particularly interested in the relationship between the encoding of anticipated (having been chosen) and offered (not yet chosen) rewards in the brain. Both types of reward are imagined, not experienced, and both can be used to influence upcoming actions. Despite these similarities, they are also somewhat conceptually distinct: offered rewards are not certain (they are contingent on choice) while anticipated rewards are generally certain. Offered rewards provide information that is used to directly drive choice, while anticipated rewards generally drive other processes, including preparation for reward receipt, savoring, and learning.

One hypothesis about the relationship between these reward types, the *simulation hypothesis*, holds that when we choose, we represent offered values and we do so by reactivating a domain-general representation of the experience of receiving the reward (Wang and Hayden, 2017; Kahnt 2010 et al., Howard et al., 2015). There is some evidence in favor of this hypothesis (Howard et al., 2015; Xie et al., 2016; Kahnt et al., 2010; Schoenbaum et al., 2003; Stalnaker et al., 2006). However, at least some data suggests that there are key qualitative differences in the way that offered and experienced rewards are represented (McNamee et al., 2015; Farovik et al., 2015; Tsujimoto et al., 2012; Wang and Hayden, 2017). These data then would predict that responses to reward in key reward regions differ depending on the context in which the reward was presented. We hypothesized that these contextual differences would show up in other domains, such as pupil size.

In order to examine the relationship between pupillary encoding of offered and anticipated values, we took advantage of an existing dataset based on two macaques performing a sequential choice task with risky options (Azab and Hayden, 2017; Azab and Hayden, 2018). We found that pupil size decreased in response to higher value offers – the opposite pattern observed for anticipated values. Consistent with this observation, pupil size following the second offer decreased less when the first was high value – a finding that is parsimoniously explained by the idea that pupils encode relative value, the key decision variable for accept-reject choices (Strait et al., 2014, Azab & Hayden 2017). Following choice, but before reward, the relationship between reward and pupil size reversed, replicating the findings of previous studies: it increased on trials in which a large reward was anticipated, and on trials in which a large reward was more likely. These findings indicate that anticipated and offered rewards are disambiguated at the level of the pupillary reward response, and suggest they are processed in at least partially distinct ways in the brain.

## METHODS

Some of the data for dorsal anterior cingulate cortex recordings were previously published (Azab and Hayden, 2017; Strait et al., 2016; Azab and Hayden, 2018; Farashahi et al., 2018; Blanchard et al, 2018). All data and analyses presented here are new.

### Surgical procedures

All procedures were approved by the University Committee on Animal Resources at the University of Rochester and were designed and conducted in compliance with the Public Health Service’s Guide for the Care and Use of Animals. Two male rhesus macaques (*Macaca mulatta*: subject B age 6; subject J age 7) served as subjects. A small prosthesis for holding the head was used. Animals were habituated to laboratory conditions and then trained to perform oculomotor tasks for liquid reward. Surgery was performed to implant a Cilux recording chamber (Crist Instruments) over the dorsal anterior cingulate cortex and subgenual anterior cingulate cortex. Position was verified by magnetic resonance imaging with the aid of the Brainsight software (Rogue Research Inc.). Animals received appropriate analgesics and antibiotics after all procedures. Throughout both behavioral and physiological recording sessions, the chamber was kept sterile with regular antibiotic washes and sealed with sterile caps. All recordings were performed during the animals’ light cycle between 8 am and 5 pm.

### Behavioral Task

Monkeys performed a two-option gambling task (Azab and Hayden, 2017; Azab and Hayden, 2018). The task was similar to one we have used previously (Strait et al., 2014; Strait et al., 2015), with two major differences. First, monkeys gambled for virtual tokens—rather than liquid—rewards. And, second, outcomes could be losses as well as wins. Our previous research confirms that subjects’ behavior is consistent with understanding of the link between colors and rewards and size and probability in this task and in ones with similar structures, including more complex foraging-like tasks – indicating that task understanding is not likely to be a limiting factor here (Azab and Hayden, 2017; Blanchard and Hayden, 2015; Blanchard et al, 2015; Sleezer et al., 2016; Ebitz and Hayden, 2016).

Two offers were presented on each trial. Each offer was represented by a rectangle 300 pixels tall and 80 pixels wide (11.35° of visual angle tall and 4.08° of visual angle wide). 20% of options were safe (100% probability of either 0 or 1 token), while the remaining 80% were gambles. Safe offers were entirely red (0 tokens) or blue (1 token). The size of each portion indicated the probability of the respective reward. Each gamble rectangle was divided horizontally into a top and bottom portion, each colored according to the token reward offered. The size of each portion indicated the probability of the respective reward. Gamble offers were thus defined by three parameters: two possible token outcomes, and probability of the top outcome (the probability of the bottom was strictly determined by the probability of the top). The top outcome was 10%, 30%, 50%, 70% or 90% likely. The possible combinations of outcomes were: +3/0, +3/-1, +3/-2, +2/+1, +2/0, +2/-1, +2/-2, +1/+1, +1/0, +1/-1, +1/-2, 0/0. Each non-safe combination was equally likely to occur.

Six initially unfilled circles arranged horizontally at the bottom of the screen indicated the number of tokens to be collected before the subject obtained a liquid reward. These circles were filled appropriately at the end of each trial, according to the outcome of that trial. When 6 or more tokens were collected, the tokens were covered with a solid rectangle while a liquid reward was delivered. Tokens beyond 6 did not carry over, nor could number of tokens fall below zero.

On each trial, one offer appeared on the left side of the screen and the other appeared on the right. Offers were separated from the fixation point by 550 pixels (27.53° of visual angle). The side of the first offer (left and right) was randomized by trial. Each offer appeared for 600 ms and was followed by a 150 ms blank period. Monkeys were free to fixate upon the offers when they appeared (and in our observations almost always did so). After the offers were presented separately, a central fixation spot appeared and the monkey fixated on it for 100 ms. Following this, both offers appeared simultaneously and the animal indicated its choice by shifting gaze to its preferred offer and maintaining fixation on it for 200 ms. Failure to maintain gaze for 200 ms did not lead to the end of the trial, but instead returned the monkey to a choice state; thus, monkeys were free to change their mind if they did so within 200 ms (although in our observations, they seldom did so). A successful 200 ms fixation was followed by a 750 ms delay, after which the gamble was resolved and a small reward (100 μL) was delivered—regardless of the outcome of the gamble—to sustain motivation. This small reward was delivered within a 300 ms window. If 6 tokens were collected, a delay of 500 ms was followed by a large liquid “jackpot” reward (300 μL) within a 300 ms window, followed by a random inter-trial interval (ITI) between 0.5 and 1.5 s. If 6 tokens were not collected, subjects proceeded immediately to the ITI.

Each gamble included at least one positive or zero-outcome, ensuring that every gamble carried the possibility of a win. This decreased the number of trivial choices presented to subjects, and maintained motivation.

Eye position was sampled at 1,000 Hz by an infrared eye-monitoring camera system (SR Research). Stimuli were controlled by a computer running Matlab (Mathworks) with Psychtoolbox (Brainard, 1997) and Eyelink Toolbox (Cornelissen et al., 2002). Visual stimuli were colored rectangles on a computer monitor placed 57 cm from the animal and centered on its eyes (Figure 1A). A standard solenoid valve controlled the duration of juice delivery. The relationship between solenoid open time and juice volume was established and confirmed before, during, and after recording.

**Figure 1.**
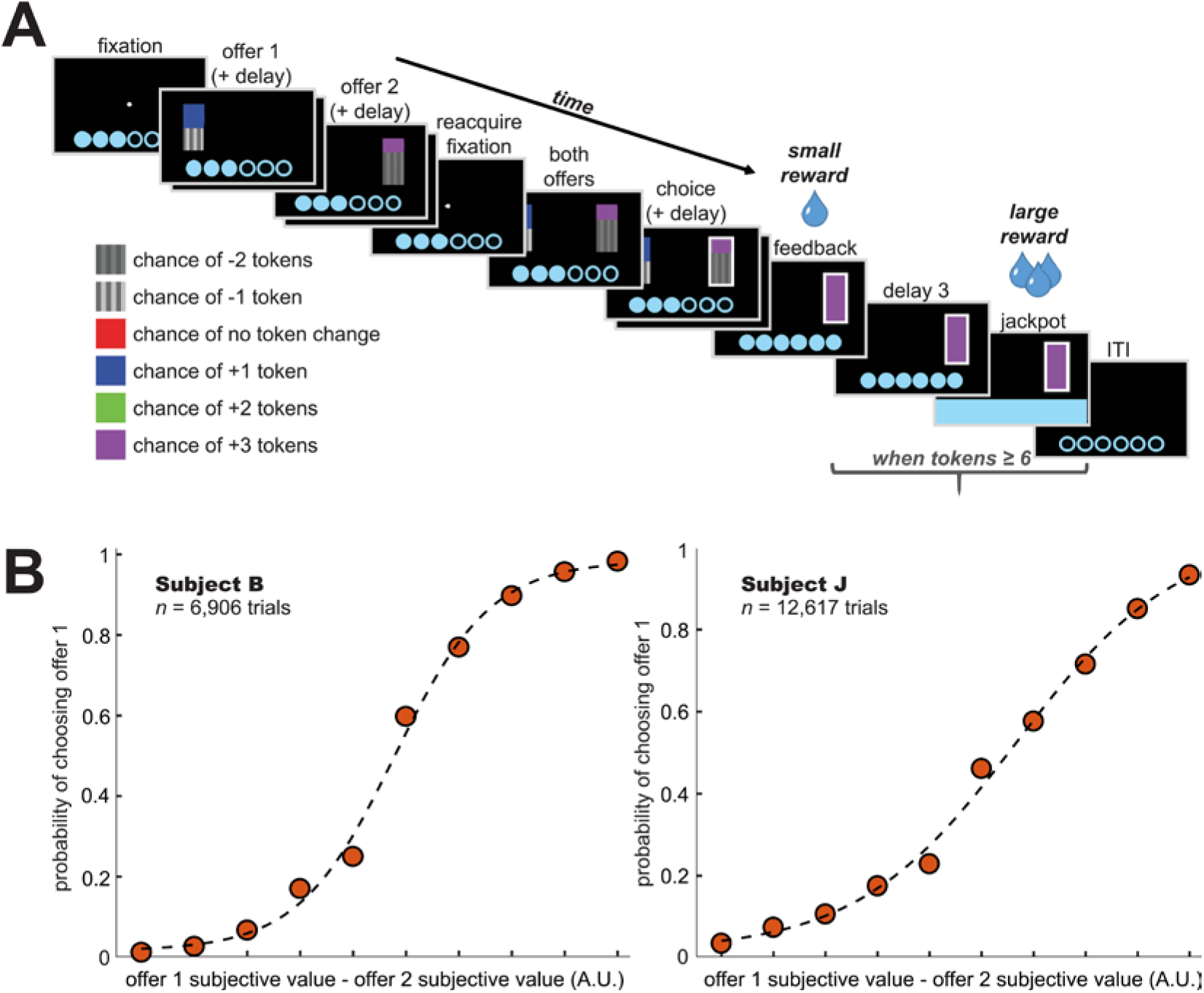
Task and Choice Behavior. (A) Token gambling task. Subjects viewed two probabilistic offers in sequence, chose between them, and then gained or lost tokens based on the result. Each offer contained two possible outcomes, represented by the color of the bars, with the probabilities of those outcomes represented the areas of those colors. (B) Choice behavior. Both subjects displayed an understanding of the task and the relative values of offers.

### Statistical Methods for Behavior

Subjective values for each gamble were estimated based on subjects’choices in each test session according to the formula:

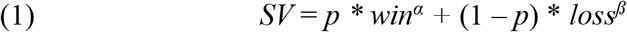

This formula comes from Yamada et al., (2013), although since our task includes both wins and losses, we fit a parameter α for wins and another parameter ß for losses. A value for α greater than 1 and a value for α less than 1 both indicate risk-seeking. Both subjects were risk-seeking on average (values of α > 1 or ß < 1 both indicate risk-seeking; subject B: average α = 1.21, average ß= 0.076; subject J: average α = 1.60, average ß = 0.022). For the remainder of this study, “value” refers to subjective value.

We fit logistic regression models of behavior to predict choice of the first vs. second offer. To ensure that subjects do, in fact, pay attention to both offers, we fit a model where the value of the first and second offers were the predictors of interest, while also including the number of tokens already accumulated, the side the first offer appears on, and the choice eventually made to explain any variance these variables might contribute to:

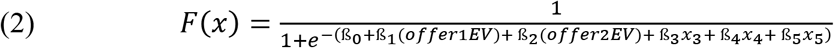

Where F(x) is the probability of choosing offer 1, x_3_ is the number of tokens, x_4_ is the side of the first offer, and x5 is the choice that was made. To determine whether subjects pay attention to all features of an offer, we use an extended model with the three variables characterizing each offer (the two possible outcomes, and the probability of the larger outcome) included as predictors, controlling for the same variables mentioned above.

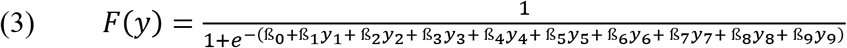

Where F(y) is again the probability of choosing offer 1, y_1_, y_2_, and y_3_ respectively are the top and bottom outcomes and probability of the top outcome for offer 1, y_4_, y_5_, and y_6_ respectively are the top and bottom outcomes and probability of the top outcome for offer 2, and then y_7_, y_8_, and y_9_ are the additional factors of token number, first offer side, and eventual choice. We fit such a model for each behavioral session, and obtain the regression weights associated with each of the variables of interest. We then test the vector of these variables across all sessions using a one-sample t-test, to determine whether they differ significantly from zero.

Trials lasting longer than the statistical ‘upper fence’—that is, the third quartile plus 1.5x the interquartile range—of trial durations were regarded as lapses and discarded. This cutoff time was calculated as the third quartile of trial length plus 1.5x the interquartile range.

### Statistical Methods for Pupil Size Analyses

We sampled subjects’ pupil diameter every 10 ms for analysis. Raw pupil data were first processed in order to remove aberrations due to blinks or measurement artifacts—outlier data points were excluded on the basis of raw size (> 99.9^th^ percentile), velocity of change (> 99^th^ percentile), and acceleration of change (> 99^th^ percentile).

Pupil sizes were then converted into z-scores on a trial-by-trial basis using the mean and standard deviation during a 500 ms normalization period, immediately preceding the start of the trial except in the instances indicated below. This length of time was chosen because it was the shortest length of the ITI, and therefore the longest normalization window that could be applied to the beginning of all trials. This normalization method was based on previously published approaches (Geng et al., 2015; Rudebeck et al., 2014), and also served the purpose of controlling for the luminance of the tokens present on the screen (we also controlled for this possibility through checking and showing no relationship, see below). During the ITI, tokens were the only object on the screen, and they remained visible on the screen throughout each trial. To analyze the effect of offer 1 value during offer 2 (figure 3), we normalized to the 500 ms preceding offer 2 onset to isolate offer-1-related pupil fluctuations and control for differing baselines. To analyze pupil effects following choice (figure 5A and 5B), we normalized pupil size to the 500 ms preceding the choice epoch, and to analyze changes in pupil size immediately following feedback (figure 5C and 5D), we normalized pupil size to the 400 ms preceding feedback appearance (the second half of the post-choice delay).

We assessed the effect of offer luminance on our results in three ways. First, we measured the screen luminance in cd/m^2^ of the color of each offer, using a Tektronix J6523-2 luminance probe, under lighting conditions identical to those of the task. The screens emitted minimal baseline luminance and there were no other sources of light during the task (figure S1). Second, we estimated the relative luminance of each offer presented to the monkey by multiplying each of the two halves’ proportion of the offer area with their respective luminance measures. We then calculated a multiple regression of the luminance and value of each offer against the mean pupil response from 150 to 350 ms following both offer 1 and offer 2 presentation on a trial by trial basis. Third, using the same time window, we performed a Pearson correlation of offer luminance and pupil response across all trials.

Offers came in several possible in a range of sizes. In our Results, ‘large’ and ‘small’ offers refer to those with subjective values greater than or less than the median offer, respectively. The onset time of offer value-related effects on pupil size was calculated by comparing the differences between the means of each trial type to shuffled data (10,000 permutations, without replacement) (Efron and Tibshirani, 1993). Similar approaches have previously been used to determine the significance of pupil size changes (de Gee et al., 2014; Nassar et al., 2012). The pupil response based on a given variable was defined as the first time bin in which the mean pupil sizes were significantly different at the threshold of α = 0.005 (two-sided permutation test). Effects with a latency of less than 250 ms were not considered, as this is approximately the shortest amount of time in which visual stimuli can induce pupil responses (Gamlin et al., 1998). Mean pupil size for large vs. small values of the first and second offer was calculated as the mean ± SEM over the 150 ms following the initial pupil response. The significance of the difference between pupil size distributions was calculated using a two-sided student’s t-test (α = 0.05).

To calculate the time course of the pupil response, we calculated the mean time of maximum pupil size difference between the two given conditions. Mean and SEM values of the maximum difference time were derived from bootstrapped data (10,000 permutations). The time window for bootstrapped data was, at a minimum, the second half of the offer epoch. In the case that significant pupil response (two-sided permutation test, p < 0.005) was observed into the delay following the offer epoch, the upper bound of the time window was either the final time bin at which a significant pupil response was observed or the onset of the next trial epoch, whichever occurred first. For the analysis of the impact of offer 1 value on pupil size following offer 2, we used a one-sided permutation test to determine the bootstrapping window in order to isolate the positive modulatory effect.

Multiple linear regression against pupil size was performed with the following regressors: offer 1 subjective value, offer 2 subjective value, number of tokens possessed, and chosen offer side. Correlations Trial-by-trial correlations with pupil size consisted were performed using the of the Pearson correlation coefficient. Correlation analyses involving offer value and pupil size were performed on a trial-by-trial basis, with pupil size calculated as the mean value during the indicated time bin. An additional regression was performed to assess the impact of pupil size during the pre-offer 2 normalization window on observed changes in pupil size following offer 2. The regressors for this analysis were offer 1 subjective value, offer 2 subjective value, and the mean pupil size during the 500 ms leading up to offer 2 onset as calculated from pupil size data that were normalized to the ITI.

To analyze the relationship between pupil size and number of tokens possessed, we excluded trials following jackpot rewards.

We performed choice probability analysis on mean pupil size during the 200 ms following offer 2 offset (the start of the pre-choice delay) from each trial. We divided trials according to whether offer 1 or offer 2 was chosen and calculated d-prime using ROC analysis (Britten et al., 1996; Britten et al., 1992). Choice probability was calculated as the area under the ROC curve. We then generated confidence intervals (α = 0.005, two-tailed) by performing similar analysis on 10,000 samples of bootstrapped data.

## RESULTS

### Choice behavior

We recorded data from two rhesus macaques in a gambling task with asynchronously presented offers (Figure 1A). Some data from this task were previously published but the data presented here are all new (Azab and Hayden, 2017; Azab and Hayden, 2018). Both subjects were familiar with the task and appeared to understand it (Figure 1B). Specifically, both subjects chose the higher value offer more than chance (subject B: 79.5% over n = 6,906 trials; subject J: 75.3% over n = 12,617 trials, p < 0.0001 in all individual sessions). Choices reflected the values of both offers according to a logistic regression model that used offer values to predict choices (see Methods, (equation 2). Both subjects showed positive regression coefficients for the first offer (one-sample t-test of coefficients for offer 1 value per session: subject B: t = 16.7; subject J: t = 27.3, both p < 0.0001) and the second offer (subject B: t = 19.7; subject J: t = 24.0, both p < 0.0001). Moreover, the values of the two possible outcomes within each offer as well as the probabilities of those outcomes all predict choices (one-sample T-test for coefficients of all 6 offer parameters: all p < 0.0001; see Methods, (equation 3).

### Increased offer value decreases pupil response

Figures 2A and 2B show the average pupil size following large and small first offers (large and small were defined relative to median offer size). During the epoch of interest (the 150 ms following onset of the response), responses were negatively correlated with the value of the first offer in both subjects (subject B: r = -0.098, R^2^= 0.010, p < 0.0001; subject J: r = -0.022, R^2^= 0.0005, p = 0.017). Immediately following onset of the offer (from 0 to 200 ms after it appeared on the screen) pupil size did not differ (this is not surprising because of the well-known slowness of pupil responses; t = -0.076, p = 0.939 for both subjects). However, following the presentation of the first offer, pupil response to large vs. small offers began to diverge rapidly. Using a sliding time window and a twosided permutation test (α = 0.005) we found that the pupil response to offer 1 value emerged at 340 ms (subject B) and 310 ms (subject J). The peak difference occurred at times 652.3 ± 0.2 ms (subject B) and 398.1 ± 0.3 ms (subject J) after offer 1 onset.

**Figure 2.**
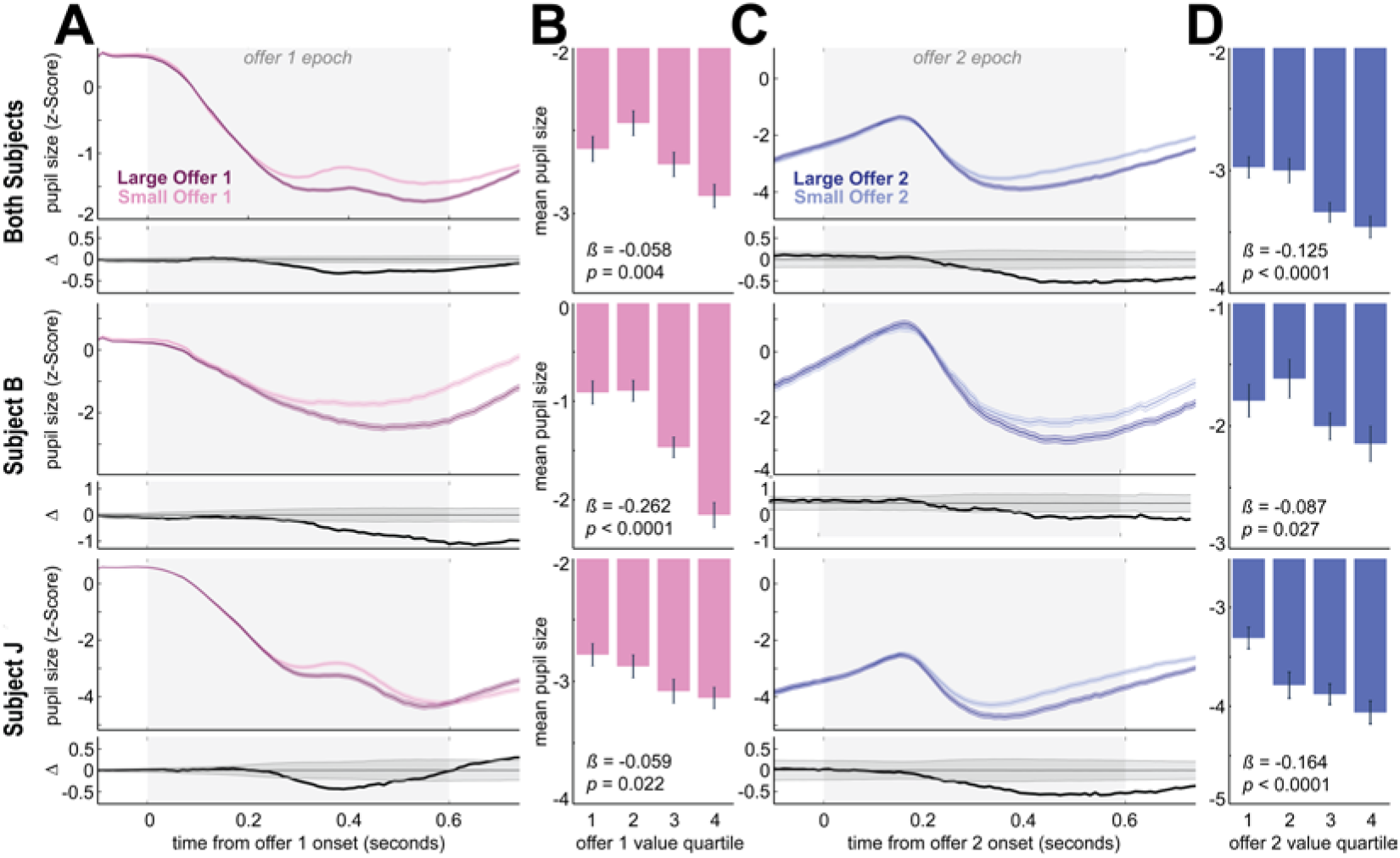
Pupil responses to offer values. (A and C) Pupil response to offer 1 (2) value during the offer 1 (2) epoch. ‘Large’ and ‘small’ refer to offers above and below the median offer value, respectively. Pupil size is normalized to the 500 ms period preceding offer 1 onset. The top panel shows the z-score (± SEM) pupil size at 10 ms increments. The bottom panel shows the difference between mean pupil size on large and small offer trials; shaded gray area shows the α = 0.05 significance threshold (two-sided permutation test). (B and D) Binned offer 1 (2) value response. Mean (± SEM) pupil size during the 150 ms following the first detected offer-related difference in pupil size. β and p values for the regression ofpupil size with offer 1 (2) value, from a multiple regression against pupil size of offer 1 and 2 value, token number, and chosen offer side, at the time of maximum difference between large and small offer responses.

At the time of peak difference, the average pupil size following large offers was significantly smaller than that following small offers (subject B: -2.006 ± 0.090 for large offers vs. -0.905 ± 0.081 for small offers, two-sided Student’s t-test, t = -8.633, p < 0.0001; subject J: -3.269 ± 0.076 for large offers vs. -2.830 ± 0.065, two-sided Student’s t-test, t = -4.334, p < 0.0001). A regression of offer 1 SV (unbinned) against average pupil size at the time of peak difference in each subject confirms this result (subject B: ß = -0.262 ± 0.033, t = -7.872, p < 0.0001; subject J: ß = -0.059 ± 0.026, t = -2.293, p = 0.022).

The same pattern was observed in the second offer epoch (Figure 2C and D). During the focal epoch, responses were negatively correlated with the value of the second offer in both subjects (subject B: r = -0.028, R^2^ = 0.001, p = 0.029; subject J: r = -0.045, R^2^ = 0.002, p < 0.0001). The difference in pupil response on the basis of offer 2 value emerged at 430 ms following the appearance of the offer for subject B and 310 ms for subject J. The peak of the difference occurred at 630.1 ± 1.2 ms (subject B) and 520.3 ± 0.6 ms (subject J). At this time, for subject B, the average size of the pupil following large offers was -2.234 ± 0.101 while the size following small offers was -1.711 ± 0.102 (these values are different, two-sided Student’s t-test; t = -3.413, p = 0.0006). For subject J, the average pupil size following large offers was -4.102 ± 0.091 while the size following small offers was -3.534 ± 0.084 (these values are different, two-sided Student’s t-test; t = -4.501, p < 0.0001). A regression of offer 2 SV (unbinned) against average pupil size at the time of peak difference in each subject confirms this result (subject B: β= -0.087 ± 0.039, t = -2.211, p = 0.027; subject J: β= -0.164 ± 0.031, t = -5.212, p < 0.0001).

### Pupil responses were not driven by variations in luminance in our task

Our offers were indicated by color, and thus varied, albeit quite modestly, in luminance. Our statistical methods were designed to eliminate confounds associated with variations in luminance (see Methods). Nonetheless, even without this control, we found no main effect of luminance in our dataset. While the average effect of offer value was strong and significant in both subjects (see above), luminance did not have significant effects (Figure S1A and B). Specifically, the luminance of offer 1 did not drive responses in either subject B (linear regression, ß = -0.002 ± 0.002, t = -0.888, p = 0.375) or in subject J (ß = - 0.002 ± 0.001, t = -1.399, p = 0.162). The luminance of offer 2 also did not drive responses in either subject B (ß = -0.001 ± 0.002, t = 0.490, p = 0.625) or in subject J (ß = 0.002 ± 0.001, t = 1.252, p = 0.211). A Pearson correlation of offer luminance and pupil response across all trials confirms this result for both offer 1 (subject B: r = -0.007, p = 0.551; subject J: r = 0.007, p = 0.406) and offer 2 (subject B: r = 0.008, p = 0.523; subject J: r = -0.015, p = 0.105).

The lack of correlation between luminance and pupil size likely reflects the relatively weak luminary effects of the small area covered by the offers (300 × 80 px on a 1024 × 768 computer monitor). It is also likely attributable in part to the stimulus colors we chose, which did not have a systematic relationship between indicated value and luminance brightness (Figure S1C).

**Figure S1.**
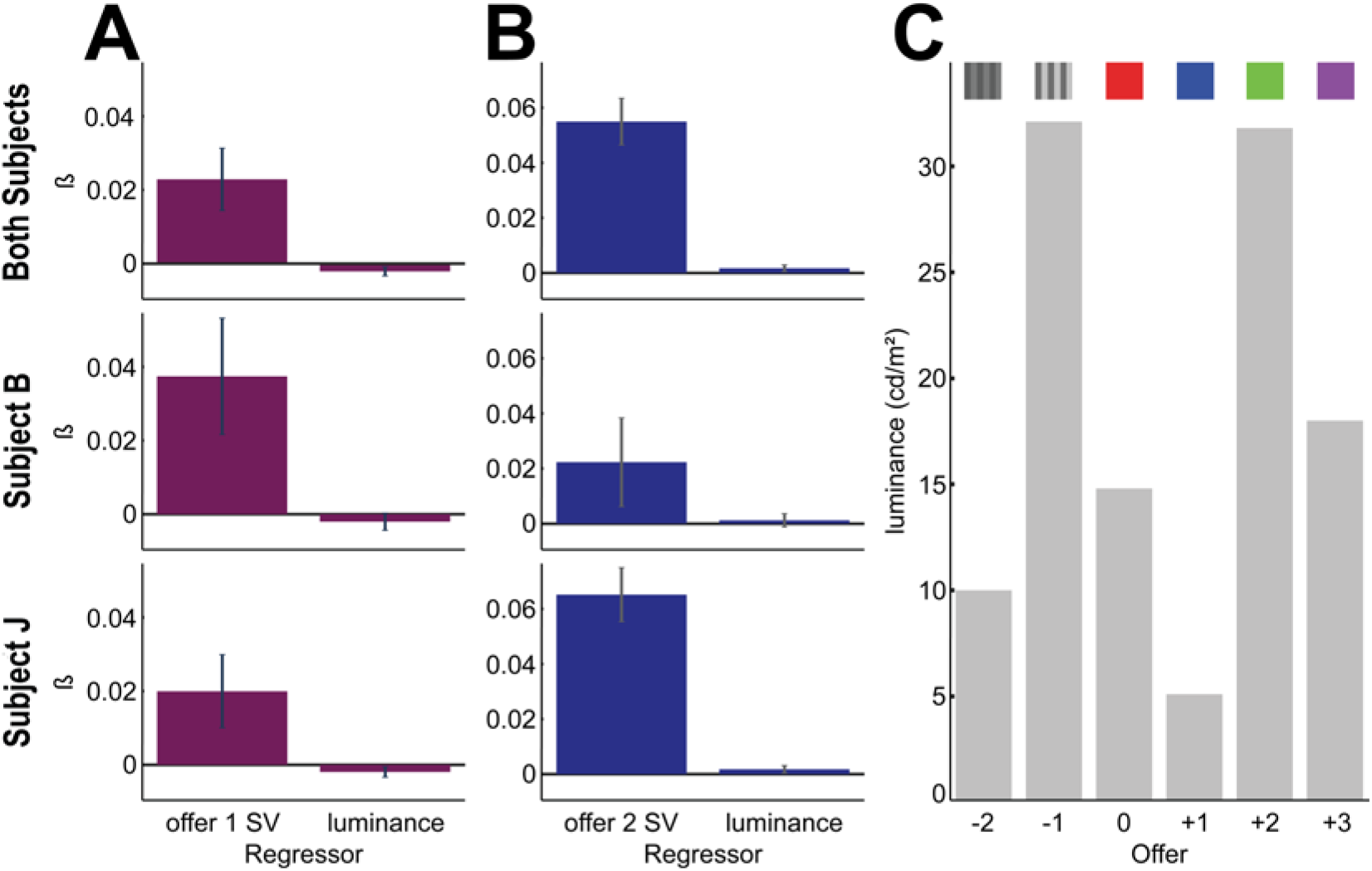
Pupil response to offer luminance. (A and B) ß values of offer 1 (2) value and luminance against pupil size during offer 1 (2). Pupil size was calculated as the mean value from the 150 ms following the onset of the first offer-related pupil size difference, the same window used for the correlations reported in this study. (C) Offer luminance. There was no systematic relationship between offer value and luminance.

### Effect of offer 1 value on the pupil response to offer 2

We next asked how offer 1 value related to the pupil response to offer 2 (Figure 3). Pupil size during the offer 2 epoch increased significantly with larger values of offer 1 (subject B: offer 1: ß= 0.228 ± 0.033, t = 7.013, p < 0.0001; subject J: offer 1: ß= 0.035 ± 0.017, t = 2.069, p = 0.039). Thus, values stored in working memory have the opposite effect of values on the screen.

**Figure 3.**
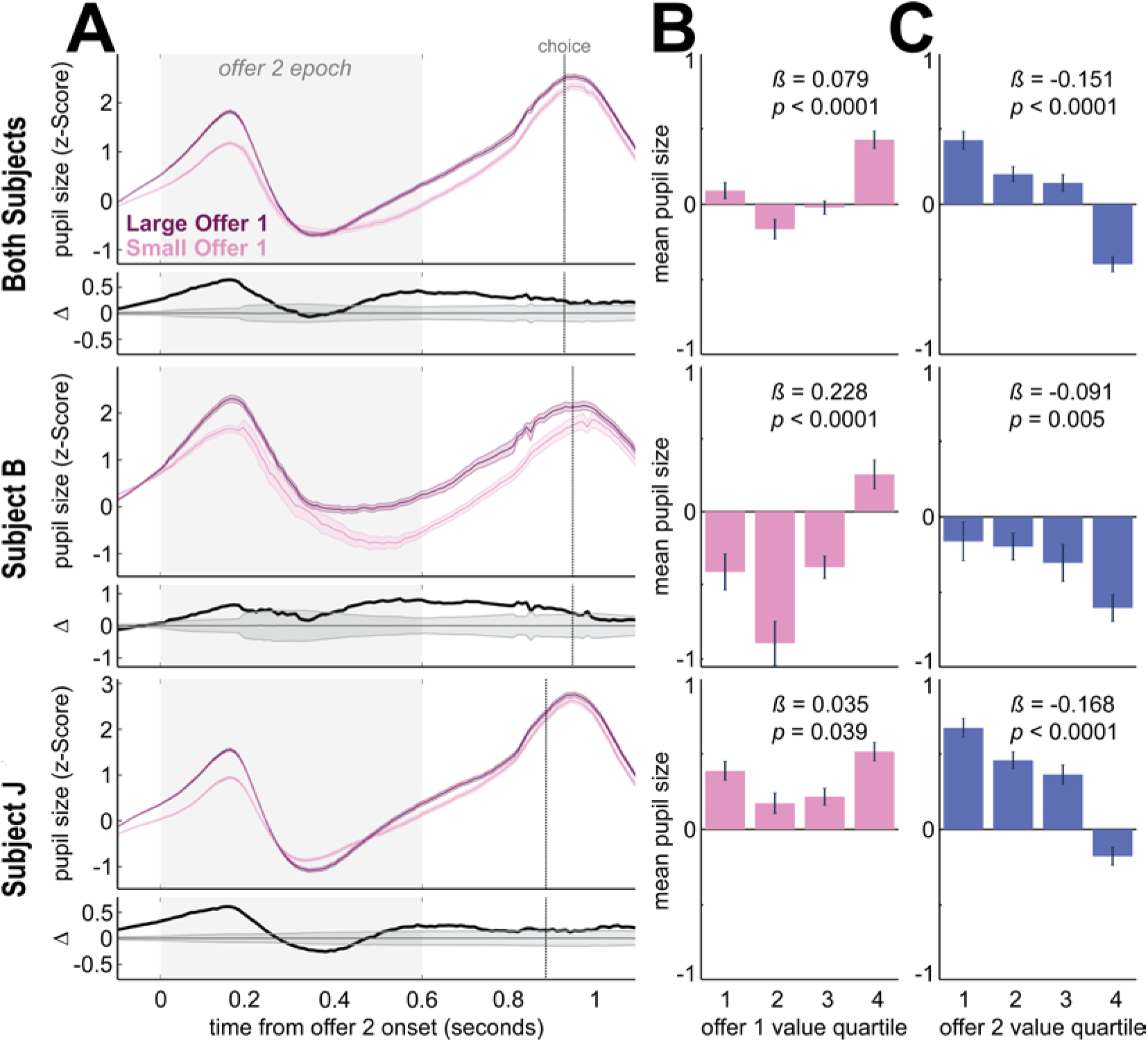
Pupil response to relative value of offer 2. (A) Pupil response to offer 1 value during and after the offer 2 epoch. ‘Large ‘ and ‘small ‘ refer to offers above and below the median offer value, respectively. Pupil size is normalized to the 500 ms period preceding offer 2 onset. The top panel shows the z-score (± SEM) pupil size at 10 ms increments. The bottom panel shows the difference between mean pupil size on large and small offer trials; shaded gray area shows the α = 0.05 significance threshold (two-sidedpermutation test). Dotted line represents the mean time of the beginning of the choice epoch for each subject, which depended on fixation time following the post-offer-2 delay. (B and C) Binned offer 1 (2) value response. Mean (± SEM) pupil size during the 150 ms following the first detected offer-2-related difference in pupil size; note that the initial 250 ms latency cutoff for stimulus-related pupil size effects (see methods). ß and p values for the regression of pupil size with offer 1 (2) value, from a multiple regression against pupil size of offer 1 and 2 value, token number, and chosen offer side, at the time of maximum difference between large and small offer 1 responses.

Specifically: for subject B, the peak offer 1-dependent difference in offer 2 response occurred at 594.9 ± 0.9 ms after offer 2 onset. At this time the average sizes of the pupil following large and small first offers were 0.164 ± 0.072 and - 0.631 ± 0.096, respectively (these values are different, two-sided Student’s t-test; t = 6.247, p < 0.0001). For subject J, the peak difference occurred at 641.1 ± 0.5 ms after offer 2 onset. At this time the average sizes of the pupil following large and small first offers were 0.511 ± 0.048 and 0.286 ± 0.046, respectively (these values are different, two-sided Student’s t-test; t = 3.316, p = 0.0009).

Since pupil size during the normalization window for this analysis (the 500 ms leading up to offer 2) includes the differential response to offer 1, it was important to account for the potential effect of this variation in the response to offer 2. To do so, we performed a regression of pupil size at the time of peak difference (measured above) against offer 1 value, offer 2 value, and the mean pupil size during the normalization window. We found that, while increased pupil size during the pre-offer2 normalization window negatively correlated with pupil size observed following offer2 (subject B: ß= -0.107 ± 0.010 t = -10.356, p < 0.0001; subject J: ß= -0.071 ± 0.005, t = -14.987, p < 0.0001), the increase in pupil size with offer 1 value and decrease with offer 2 value both remained significant (subject B: offer 1: ß= 0.200 ± 0.023, t = 8.900, p < 0.0001; offer2: ß= -0.053 ± 0.022, t = -2.392, p = 0.017; subject J: offer 1: ß= 0.037 ± 0.014, t = 2.552, p = 0.011; offer2: ß= -0.1529 ± 0.014, t = -10.684, p < 0.0001).

### Pupil size predicts choice and reflects the value of the chosen offer more strongly than the value of the unchosen offer

Following the presentation of the second offer, pupil size steadily increased leading up to the choice epoch (Figure 4). During the pre-choice delay, when no offer stimuli were on the screen (0 ms to 200 ms following offer 2), pupil size was correlated with the value of the chosen offer in the two subjects together (r = -0.019, p = 0.010), and was significant in one subject and close, but not statistically significant, in the other (subject B: r = -0.024, p = 0.051; subject J: r = -0.023, p = 0.009). It was not correlated, however, with the unchosen offer in either subject (subject B: r = -0.008, p = 0.540; subject J: r = -0.010, p = 0.260), or in the two subjects averaged together (r = -0.008, p = 0.294). These findings are consistent with the idea that following presumed covert choice subjects attend the value of the chosen offer more than the value of the unchosen offer (Hayden & Moreno-Bote, 2017).

**Figure 4.**
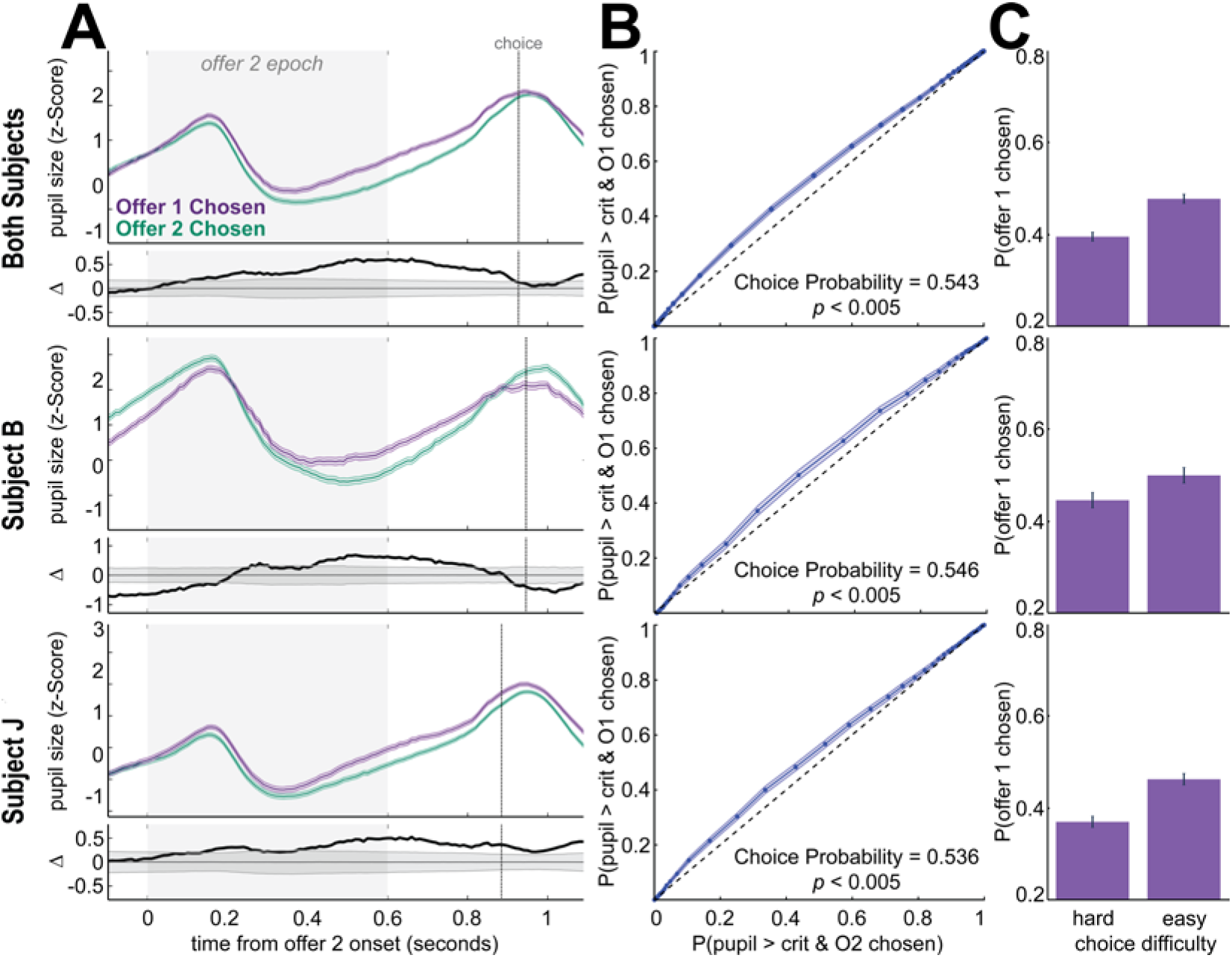
Pupil size by chosen offer and choice heuristic. (A) Pupil size by chosen offer during and after the offer 2 epoch. Pupil size differed on the basis of which offer was ultimately chosen. Pupil size is normalized to the 500 ms period preceding offer 1 onset. Dotted line represents the mean time of the beginning of the choice epoch for each subject, which depended on fixation time following the post-offer-2 delay. (B) Pupil size predicts choice. Analysis window was the 200 ms following the offer 2 epoch. Choice probability is the area under the ROC curve, indicating the probability that an ideal observer could predict the chosen offer from only the mean pupil size during the analysis window. (C) Choice heuristic. ‘Hard’ and ‘easy’ trials refer to trials in which the two offers were below or above the median difference in offer values, respectively. Both subjects chose offer 1 significantly more often on ‘easy’ trials than on ‘hard’ trials (p < 0.05).

### Following choice, pupil size increases with anticipated value

In the delay following choice, while subjects awaited feedback on their gamble, pupil size was higher when the jackpot reward was within reach. Specifically, we compared pupil size on trials when subjects possessed 3 or more tokens to trials when they possessed fewer (two-sided Student’s t-test; Subject B: t = -4.602, p < 0.0001; subject J: t = -13.834, p < 0.0001). This effect is not dependent on binning: regressing pupil size by number of tokens demonstrated a significant positive relationship (Figure 5B; subject B: r = 0.954, p = 0.003; subject J: r = 0.866, p = 0.026).

**Figure 5.**
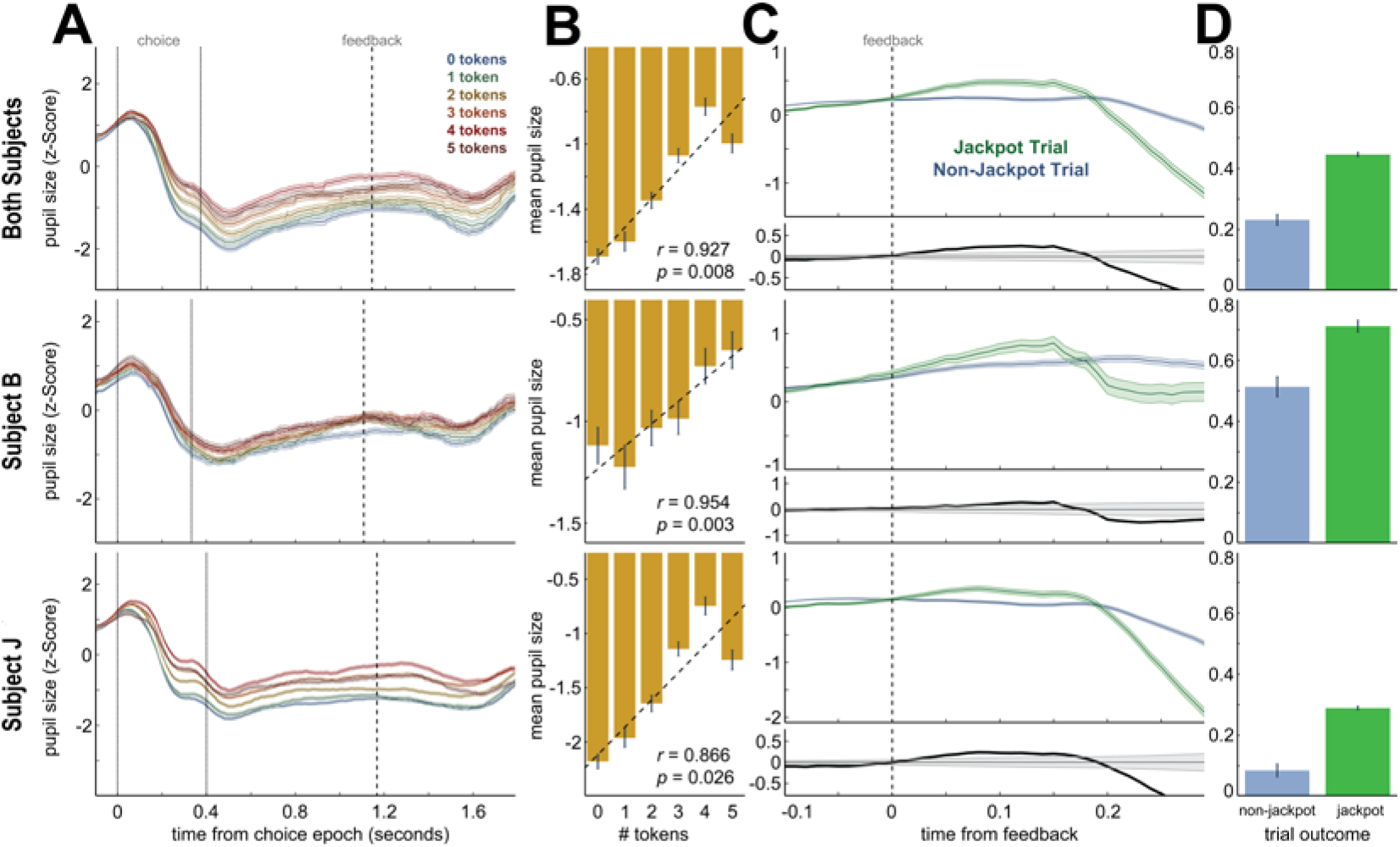
Pupil response to anticipated value. (A) Pupil size by number of tokens possessed during the trial. Pupil size tended to increase on trials in which the subject possessed more tokens, and thus the jackpot reward was more likely. Pupil size in 5A and 5B is normalized to the 500 ms period preceding choice epoch onset, and for figures 5C and 5D to the 400 ms period preceding feedback delivery. The lines indicate the z-score (± SEM) pupil size at 10 ms increments. The dotted lines represent beginning and mean end time of the choice epoch, and the dashed line represents the mean time of feedback presentation. (B) Mean pupil size from 400 to 500 ms following choice epoch onset, binned by number of tokens possessed. Pupil size increased with number of tokens. Dashed line indicates a linear regression of the mean bin values; r indicates the Pearson correlation coefficient and p the significance level. (C) Pupil size following feedback on jackpot vs. non-jackpot trials. Pupil size transiently dilated on trials in which subjects expected to receive the jackpot primary reward. (D) Mean (± SEM) pupil size following feedback on jackpot vs. non-jackpot trials. Pupil size was averaged over the 200 ms following feedback, a time period before the actual feedback information could be integrated into pupil size.

In our task, there was a delay following feedback and before the reward itself. A transient pupillary dilation coincided with the delivery of feedback on jackpot trials, demonstrating that subjects anticipated the large primary reward itself (Figure 5C). Pupil size was larger during the period immediately following feedback on jackpot trials (Figure 5D; subject B: 0.712 ± 0.020; subject J: 0.289 ± 0.006) than on non-jackpot trials (subject B: 0.513 ± 0.033; subject J: 0.085 ± 0.021). These effects were significant in both subjects (2-sided Student’s t-test; subject B: t = 2.599, p = 0.009; subject J: t = 4.060, p < 0.0001). Note that the appearance of feedback itself on jackpot trials (a blue bar across the bottom of the screen; see figure 1A) resulted in pupillary constriction, but the anticipatory dilation occurred during the ~200 ms immediately following feedback, before any seen information could be expected to be expressed in pupil size.

### A heuristic bias in choice

We observed a novel heuristic choice bias in this dataset. The effect builds on a recency bias previously observed in macaques and extended here (Blanchard, Wilke et al., 2014; Blanchard, Wolfe, et al., 2014). Specifically, subjects chose offer 2 slightly more often than offer 1 (subject B: 52.8% ± 1.19%, subject J: 58.3% ± 0.86%, binomial proportion 95% confidence intervals, binomial test, p < 0.0001 for both subjects). The novel finding is that subjects were more accurate (i.e. more likely to choose the EV-maximizing option) when they chose offer 1 (subject B: 80.96% ± 1.37% vs. 74.83% ± 1.43%; subject J: 77.93% ± 1.13% vs. 70.22% ± 1.05%, binomial proportion 95% confidence intervals, Fisher’s exact test, p < 0.0001 for both subjects). This bias can be explained by a sequential, accept-reject choice process: if subjects attend to and decide on one offer at a time, then at the time of choice offer 2 will tend to be the attended offer and therefore accepted more often by default. Supporting this interpretation, the bias towards offer 2 was more pronounced when decisions were difficult--that is, when options were more similar in SV (Figure 4C, Fisher’s exact test, p < 0.0001 for both subjects).

Further supporting this idea, the strength of the bias decreased with increasing number of tokens. In our task, all trials were followed by the same amount of primary reward, except for trials on which monkeys successfully accumulated six tokens and subsequently received a large, ‘jackpot’ primary reward. For this reason, token number, which was displayed throughout the trial (including during the intertrial interval), provided a running measure of proximity to this large reward. When subjects had 5 tokens, 1 token away from the large primary reward, they chose offer 1 49.63% ± 3.45% of the time; when subjects had fewer than 5 tokens, they chose offer 1 only 43.10% ± 1.27% of the time (Fisher’s exact test, subject B: p = 0.038; subject J: p < 0.0001). Response times also decreased when subjects possessed 5 tokens (384.2 ± 2.6 ms) vs. when they possessed fewer than 5 tokens (420.9 ± 1.4 ms; t = -8.901, p < 0.0001 for both subjects). These data suggest that offer 2, as the putative attended offer during the choice epoch, is processed more easily than offer 1 except under highly motivated conditions.

## DISCUSSION

We examined the relationship between offered and anticipated values on pupil size in rhesus macaques performing a sequential choice task with asynchronous offer presentation. Larger offered values for both the first and second offers led to stronger pupillary constrictions. The value of the second offer was encoded relative to the value of the first and not absolutely. Pupil size thus negatively tracks the relative value of the presumed attended offer. Immediately before choice, when neither offer was visible, pupil size correlated with the value of the chosen, but not unchosen offer. Following choice, pupil size was larger on trials with greater token count, which is correlated with higher base trial value. Furthermore, pupil dilation coincided with feedback on jackpot trials, when a large reward was expected. Our results confirm previous findings on anticipated value and pupil size and extend them to new contexts. We also show a relationship between offered value and pupil size, one that is the opposite of the previously published relationship between anticipated value and pupil size.

Our results indicate a clear dissociation in the effect that offered and anticipated rewards have on pupil size, and thus suggest they are not processed in the same way. These results therefore argue against the strongest versions of the *simulation hypothesis* – the idea that the way we represent offered rewards is to reactivate states associated with anticipating (and in some cases, receiving) the reward (Kahnt et al., 2010; Schoenbaum et al., 2003; Stalnaker et al., 2006; Wang and Hayden, 2017; Xie et al., 2016). Instead, they are consistent with the idea that the representation of offered rewards has elements that are qualitatively different from those of anticipated rewards, leading to the difference in the way they are reflected in pupil size. This distinction is also reflected in the way offered and anticipated rewards are encoded in neural responses (Farovik et al., 2015; McNamee et al., 2015; Tsujimoto et al., 2012; Wang and Hayden, 2017). Note that our results do not argue against a weaker version of the simulation hypothesis, in which offers involve partial reactivation of response patterns associated with receipt but also activate other orthogonal response patterns. Other results from our lab provide neuronal evidence in favor of that hypothesis (Wang and Hayden, 2017).

We have previously argued that it can be helpful to take a foraging perspective to understand economic choice (Cisek, 2012; Hayden, 2018; Pearson et al., 2014). From this perspective, decision-makers consider one option at a time, evaluate the value of accepting it relative to the value of rejecting it, and then accept it if the value is above some threshold (Freidin et al., 2009; Kacelnik et al., 2011). This finding is echoed in neural responses (Krajbich et al., 2010; Rich and Wallis, 2016) and, here, in pupil responses. Across the two offers, the size of the pupil is correlated with the size of the attended value. Indeed, the decrease in pupil size with increasing offered value may indicate attention to the presented offer. The converse would then be true for the case of offer 2 on trials in which offer 1 was highly valuable. That is, if the subject “accepts” a highly valuable first offer, he may then pay less attention to the second and be less focused in the prechoice delay, leading to the larger pupil size that we observed during those epochs. This idea has an echo in classic idea of memory-guided decision-making, in which attention to a memorandum alters the responsiveness of high-level association neurons to subsequent probes, allowing for fast feed-forward decisionmaking (Miller and Desimone, 1993; Miller et al., 1991; Mirabella et al., 2007; Hayden and Gallant, 2013; Lui and Pasternak, 2011; Machens et al., 2005).

The observation of a heuristic bias in our subjects toward choosing offer 2 is consistent with this hypothesis. The second offer, being presented right before the choice epoch, would naturally tend to be the attended offer at the time of choice. The subject would thus be predisposed to accept it. This may have resulted in the generally quicker choices and preference on difficult trials that we observed toward offer 2 in both subjects. On the other hand, when offer 2 does not meet the threshold for acceptance, a subject must engage in the more cognitively demanding process of shifting attention to and evaluating offer 1. For marginal decisions, this may have only been ‘worth it’ to the subjects when they had 5 tokens and were thus close to receiving a large primary reward—the only condition under which the bias toward offer 2 disappeared. Along these same lines, but assuming less of a link between the value-based decision making system and pupil-linked arousal, is the possibility that the higher pupil size in response to low-value offers indicate that the subject is resisting the natural inclination to accept what is currently available.

An advantage of viewing decision making through the lens of foraging is that it provides a new perspective on the fundamental meaning of value, one of the important philosophical problems of neuroeconomics (Hunt and Hayden, 2017; Levy and Glimcher, 2012; O’Doherty, 2011; Schultz, 2008; Wallis and Rich, 2011). Specifically, it suggests that value is not a single entity, but a convenient name for a variety of constituent cognitive processes. These processes are not necessarily highly correlated; indeed, they can, as in the case of offered and anticipated values, have opposing effects. They are also likely to be broadly distributed throughout the brain, rather than bound to a particular population or area. It is therefore no surprise that their distinct traces show up in such a global indicator of brain state as pupil size.

## Statement on the Welfare of Animals

All procedures performed in this study involving animals were in accordance with the ethical standards of the University of Rochester.

## Acknowledgments

This work is supported by a CAREER award from NSF (BCS1253576) and a R01 from NIH (DA038615) to BYH. We thank Meghan Castagno, Marc Mancarella and Caleb Strait for assistance with data collection, and the rest of the Hayden lab for valuable discussions.

